# CA1 reinstatement and novelty response independently support recognition memory

**DOI:** 10.64898/2026.09.14.751602

**Authors:** Jörn A. Quent, Mengya Zhang, Deniz Vatansever

## Abstract

One of the central tensions in memory research is whether consistency or differentiation across events is beneficial for remembering. While theoretical models posit that the hippocampal subfield CA1 subserves both types of computations, it remains unclear how it can simultaneously express both in service of recognition memory. Here, we contrasted the two prominent functions of CA1: reinstatement (consistency) and novelty processing (differentiation) to address this gap. To this end, we analysed the Natural Scenes Dataset featuring eight participants who completed a continuous recognition memory task spanning 41 separate days with 213,000 trials in total. We calculated reinstatement and novelty indices for each image and then examined the degree to which these reflected successful memory encoding. We found that CA1 showed evidence for encoding-related reinstatement as well as for novelty processing even after controlling for each other. Reinstatement was specific to retrieving the initial representation during the first retrieval attempt and did not extend to other forms of reinstatement. Exploratory analyses additionally identified reinstatement in CA3, which exhibited different response profiles compared to CA1. Our work thus highlights that computations of consistency as well as differentiation can be orthogonal signals contributing independently to memory success and can simultaneously co-exist within the same region of the hippocampus.

**Significance statement:** One of the central tensions in memory research is whether consistency, measured by reinstatement or differentiation, measured by activation difference across repeated events benefits remembering. While both have been associated with the hippocampal subfield CA1, it remains unclear how they can simultaneously support recognition memory. Here, we leveraged 7T precision fMRI in a densely-sampled continuous recognition task and demonstrated that reinstatement of the first representation and the novelty response co-exist in CA1. Critically, both independently predict subsequent memory success, suggesting computations of consistency and differentiation are orthogonal signals. We additionally identified subfield CA3 in which reinstatement, but not novelty, is predictive of memory, further highlighting the unique role of CA1 as a convergent hub of multiple hippocampal functions.

## Introduction

Our ability to remember relies on encoding new information to enable subsequent retrieval. On the one hand, repeated events can trigger reinstatement with beneficial effects for memory (Ritchey et al., 2013; Xue, Dong, et al., 2010), suggesting consistency across repetitions is advantageous. On the other hand, many brain regions also distinguish between novel and familiar stimuli (Kim, 2013). This activation difference (i.e. novelty response) – and thus differentiation – has also been associated with successful memory formation (Tulving et al., 1996; Tulving & Kroll, 1995). Yet, it is unclear how both consistency and differentiation can produce memory benefits simultaneously.

A similar debate revolves around the hippocampus – the central memory region, whose computations support consistency (Horner et al., 2015; Silva et al., 2026) as well as differentiation (Bakker et al., 2008; Norman, 2010; Yassa & Stark, 2011). Briefly, within the hippocampus, CA1 in this circuitry receives direct projections from the entorhinal cortex but also indirect projections relayed via DG and CA3 (Hasselmo & Wyble, 1997). This places CA1 in a unique position to serve as a comparator (Bein & Davachi, 2026).

For instance, CA1 showed higher pattern similarity for objects sharing the same virtual context, while CA23/DG represented these as more dissimilar (Dimsdale-Zucker et al., 2018). The hippocampus (Libby et al., 2019) but especially CA1 is pivotal to integrating and generalising information, allowing associative inference (Schlichting et al., 2014) as well as hippocampal reinstatement (Tompary et al., 2016). The re-expression of neural representations of the initial experience in CA1 predicted for example the precision of temporal judgements (Zou et al., 2023). Together, these findings indicate that hippocampal subfields may underpin dissociable processes with CA1 being particularly important for reinstatement.

The tension between consistency and differentiation also cuts across the question regarding which univariate activation patterns are signatures of successful memory. The hippocampus often shows higher activation for novel relative to familiar information (Kim, 2013; Kumaran & Maguire, 2009; Quent et al., 2021; Quent, Song, et al., 2026). It has long been hypothesised that this reflects differential encoding activity (Kim, 2013; Kirchhoff et al., 2000; Tulving et al., 1996; Tulving & Kroll, 1995). Novelty processing and its effects on behaviour are likewise strongly associated with CA1 (Chen et al., 2015; Duncan et al., 2012; Maass et al., 2014). For example, recent work demonstrated that CA1 alongside the subiculum plays a role in novelty detection and recognition memory (Quent, Zhuang, et al., 2026). Importantly, there are several lines of evidence to suggest that the strength of the novelty response is a signature of successful memory formation (Ben-Yakov et al., 2014; Maass et al., 2014; Turk-Browne et al., 2006), at the same time there is also support for consistent encoding responses across the brain being beneficial (Manelis et al., 2013; Wagner et al., 2000; Xue et al., 2011; Xue, Mei, et al., 2010). This therefore raises the question how the strength of the novelty response in CA1 relates to memory formation and how it can be squared with its role in reinstatement.

The relationship between novelty and reinstatement in humans therefore deserves further study. Previous work on the cortex suggests that reinstatement mostly supports recognition memory, while the novelty response mostly supports repetition priming (Ward et al., 2013). This suggests dissociable roles of novelty processing and reinstatement in memory, but it remains unclear whether this generalises to the hippocampus and to the division of labour among its subfields. One possible explanation for this apparent gap is that evidence for hippocampal reinstatement has been surprisingly elusive (Ritchey et al., 2013; Staresina et al., 2012; Wing et al., 2015). A potential reason is that most studies average across subfields despite their diverse mnemonic contributions. It has also been suggested that goals flexibly modulate hippocampal representations, shifting processing between integration and differentiation (Brunec et al., 2020). However, recent evidence points out that even seemingly minor methodological differences play a larger role in shaping hippocampal responses than previously appreciated (Yuksel et al., 2026). This illustrates the need for further work on how the hippocampus and specifically CA1 can support reinstatement as well as novelty processing in service of memory.

## Methods

### Natural scenes dataset

Given the elusiveness of hippocampal reinstatement, we decided to analyse the Natural Scenes Dataset (NSD; Allen et al., 2022), which is a large-scale 7T-fMRI dataset featuring a densely sampled continuous recognition memory task. We chose this dataset because i) during a recognition memory task the perceptual input remains constant; only mnemonic processes differ. ii) lags between repetitions in NSD span from minutes to months, allowing us to probe whether hippocampal novelty processing and reinstatement are time-limited, greatly improving generalisability. iii) NSD also affords the necessary statistical power – with 213,000 individual trials in total – to find potentially subtle effects. We hypothesised that CA1 is the primary candidate to process novelty (Chen et al., 2015; Duncan et al., 2012; Maass et al., 2014) as well as to express consistent representations across repetitions in the form of reinstatement (Tompary et al., 2016; Zou et al., 2023).

A detailed description of the NSD (http://naturalscenesdataset.org) is provided elsewhere (Allen et al., 2022). In short, NSD contains measurements of fMRI responses from eight participants who each viewed between 9,000 and 10,000 distinct colour natural scenes (22,000–30,000 trials) over the course of 30–41 scan sessions. After the completion of the scanning sessions, participants completed an additional final memory task, which was not analysed here (see below for reasons). Scanning was conducted at 7T using whole-brain gradient-echo EPI at 1.8 mm resolution and 1.6 s repetition time. Images were taken from the Microsoft Common Objects in Context (COCO) database (Lin et al., 2014) and cropped to be square with a size of 8.4° x 8.4°. Images were presented for 3 s with 1 s gaps between images. Participants fixated centrally and performed a long-term continuous recognition task on the images (Figure 1A). The fMRI data were pre-processed by performing one temporal interpolation (to correct for slice time differences) and one spatial interpolation (to correct for head motion). A general linear model was then used to estimate single-trial beta weights (Prince et al., 2022). Since our analyses focussed on potential changes across repeated presentations of the same stimulus, in line with similar investigations (Zou et al., 2023), we used TYPEB betas (i.e. betas_fithrf) at the 1.8 mm preparation, which were derived by optimally selecting haemodynamic response functions from a library of 20 candidates. These single-trial betas represent signal percent change multiplied by 300 to save disk space. No z-scoring at the level of the raw betas was applied.

**Figure 1.**
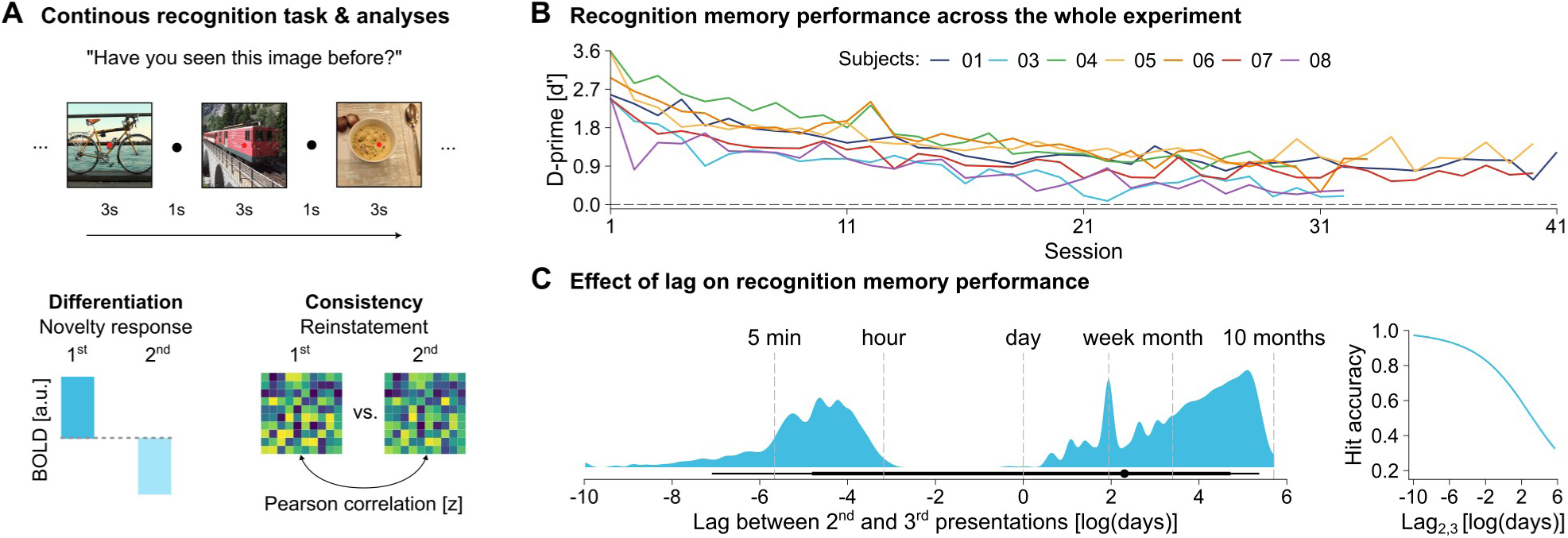
(A) Illustration of the continuous recognition task. Participants viewed a series of naturalistic scenes. On each trial, they had to indicate whether they have seen this image before in the experiment. **(B) Recognition performance over the course of the experiment.** Lines represent memory performance (d-prime) for each individual subject across sessions. **(C) Effect of lag on recognition performance.** The distribution of log-transformed lags in days is shown in a density plot with the median and 66 % and 95 % CI illustrated below. The relationship between lag and subsequent memory is illustrated via marginal predictions of a hierarchical logistic regression model. COCO photos from https://flickr.com: bicycle (user: russteaches), train (user: kecko), bowl of food (user: find Your Feet), CC BY 2.0 (https://creativecommons.org/licenses/by/2.0/).

Informed written consent was obtained from all participants. The experimental protocol of the NSD project was approved by the University of Minnesota institutional review board (Date: August 3, 2018, IRB ID: 1508M77463, Submission ID: MODCR00001243).

### Hippocampal segmentation

We segmented the hippocampus and generated individual surfaces using HippUnfold (DeKraker et al., 2020) via a singularity container (v1.5.2). HippUnfold was applied to the 0.5 mm preparation of the T1 image (“T1_0pt5_masked.nii.gz”) with output density set to 0.5 mm (see https://github.com/cognizelab/NSD_HippUnfold for the full pipeline). Using the pre-computed transformations (“func1pt8-to-anat0pt8.nii.gz”) and adapted code from NSD’s utility functions (https://github.com/cvnlab/nsdcode), we transformed the functional data to align it with the anatomical space. Next we mapped the single-trial beta estimates to the hippocampal surfaces with ribbon constraint using the inner, midthickness as well as outer surfaces. Finally, the data were downsampled to 2 mm resolution and saved as “.h5” files for simplified data handling. This pipeline used Connectome Workbench’s wb_command utilities, HippoMaps (DeKraker et al., 2025) as well as custom Python and shell code made fully available with this publication. In addition to visual inspection, hippocampal segmentation success was evaluated based on Dice coefficients estimated by the HippUnfold pipeline. In line with recommendations, coefficients exceeded 0.7 for all segmentations apart from sub-02’s left hippocampus, which was only 0.66. We therefore excluded this participant from all analyses. Based on the final sample size (N = 7), Dice coefficients averaged across hemispheres ranged from 0.80 to 0.83 (mean = 0.82, SD = 0.013).

Our primary region of interest (ROI) was the bilateral CA1, which along with the other subfields, was defined based on the multihist7 atlas (DeKraker et al., 2023). In exploratory analyses, we also examined the presence of hippocampal reinstatement in the subiculum, CA2, CA3 and DG to probe the specificity of potential effects in CA1 (Figure 2A). Due to its size and the limited resolution of the data, CA4 was not included in these analyses.

**Figure 2.**
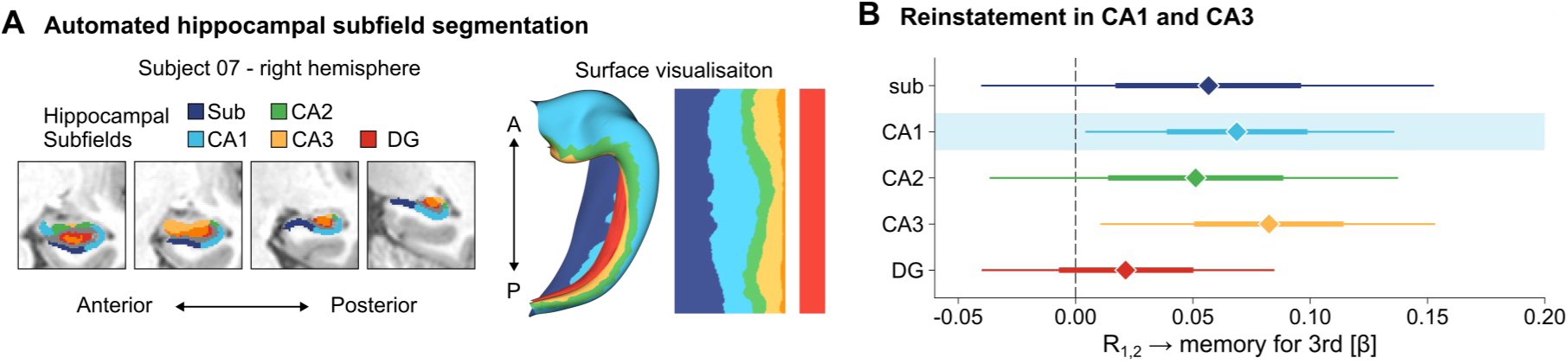
(A) Automated hippocampal segmentation. Hippocampal subfields and surfaces were segmented and generated using HippUnfold. Segmentation success is illustrated for sub-07 in four representative slices of their T1w image with segmented subfields. Additionally, we show the canonical hippocampal surface to illustrate the locations of the subfields. **(B) Hippocampal reinstatement in CA1 and CA3.** Posterior distributions of the fixed effects are summarised via their median (centre diamond) and the 66% and 95% CI. The primary ROI, CA1, is highlighted.

### Statistical analyses

First, we examined whether reinstatement of the initial experience in the region is a signature of successful memory formation. Conditional on finding such evidence, we further sought to understand how these consistent representations can be squared with the activity difference of the novelty response (Figure 1A). Prior to any statistical analysis involving behaviour, trials with faster responses than 300 msec or trials with no responses were excluded. All details on specific software and packages used for this publication can be found in the accompanying code repositories.

### Bayesian hierarchical regression analyses

To test our hypotheses, we used several Bayesian hierarchical regression models fit using the R package brms (Bürkner, 2017). The same procedure was used for all models, all of which included full random effects (slopes and intercepts) for individuals. Because of the small number of units (i.e. participants; N = 7), we kept the random effect structures as simple as possible without artificially inflating our fixed effect precision. Note, including correlations between random effects as well as dropping the random slopes all together inflate the estimation of precision (i.e. lower the standard error). To increase the robustness of our statistical inference, we choose the most rigorous and conservative approach by including random slopes while dropping the correlations between random effects (Barr et al., 2013) even at the cost of enlarging our credibility intervals (CI).

Models were fit using five chains each with 6,000 iterations, half of which were used for warm-up. All models finished with an R-hat of 1 with no divergent transitions. Predictor variables were all scaled to have a mean of 0 and a SD of 0.5. For Gaussian regression models, the independent variable was scaled to have a mean of 0 and a SD of 1. For the trial-level analyses, priors for neural effects were chosen in line with effect sizes reported in prior work (Quent, Zhuang, et al., 2026), where the univariate β for the novelty response was between 0.03 and 0.085 on the standardised scale. The maximum β for a logistic regression model was 0.32 for similar comparisons. We therefore chose the width of our priors so that 50% of the mass just included these values for our neural indices (e.g. N(0, 0.47) for logistic models). For all models, we report the regression coefficients and their corresponding 95% credibility intervals (CI). If the CI included zero, an association was interpreted as not being present/robust. The regression coefficients for the interactions were only reported in cases where their CI did not include zero.

Note that we also investigated whether any of our neural measures predicted performance in the final memory task, but none of those predictions were robust. This can be partly attributed to the reduced power because the final memory test had an order of magnitude fewer trials and/or to the criteria used to select trials (Zou et al., 2023). These analyses are therefore not reported here.

### Behavioural analyses

As shown in previous work, the NSD affords a high level of data quality and behavioural performance, which has been extensively reported elsewhere (Allen et al., 2022; Koslov et al., 2024; Vanasse et al., 2022; Zou et al., 2023). To avoid repetition, we restricted our analyses to immediately relevant issues. First, we demonstrated that average memory performance (measured by d-prime) remained at moderate levels of discrimination throughout most of the experiment. To quantify how performance changed over the course of the experiment, we modelled session-level averages of d-prime as a function of the session number. We decided against excluding later sessions to increase the generalisability of our investigation and to specifically investigate how variations in encoding potentially protect against forgetting and retrieval failure. Furthermore, we examined the role of lag between presentations in successful recognition judgements. Since our main interest was to explain memory for the 3^rd^ time, we first modelled trial-wise memory performance as a function of the log-transformed lag between the 2^nd^ and the 3^rd^ encounter as measured in days, using a logistic regression model. Additionally, we modelled memory for the 2^nd^ presentation to illustrate how the effect of lag on memory compares with the effects of lag on novelty processing and reinstatement.

### Reinstatement analyses

Three presentations of a stimulus allow calculations of several types of reinstatement. The main type of reinstatement we were interested in here was the reactivation of the representation of the initial experience when it was explicitly prompted for the first time (i.e. the 2^nd^ presentation), which previous investigations have found to be strongly associated with subsequent memory (Heinbockel et al., 2024; Zou et al., 2023). This choice was made because we were interested in reinstatement’s contribution to memory formation. This reinstatement is denoted as R_1,2_ because it was calculated by correlating the activation vectors of the 1^st^ and 2^nd^ encounter within a subfield. Our main interest was to quantify the extent to which this form of reinstatement relates to subsequent memory (i.e. for the 3^rd^ encounter).

Additional exploratory analyses investigated how other forms of reinstatement (e.g. R_1,3_ and R_2,3_) related to memory. In addition to calculating reinstatement for pairs of encounters, we calculated R_avg,3_ to investigate the possibility that during the 3^rd^ encounter participants actually reinstated an integrated representation of the 1^st^ and 2^nd^ presentations, which was achieved by summing the activation vectors prior to calculating the pattern correlation. All Pearson correlations were Fisher z-transformed before statistical analysis.

### Tests of item and context-specificity

After establishing that there was evidence for hippocampal reinstatement, we examined its nature. A positive relationship between R_1,2_ and subsequent memory can arise via several mechanisms. One possibility is that during the 2^nd^ encounter participants reinstate the specific representation that is exclusively associated with a particular item, which is predicted by the standard reactivation account. However, it is also possible that a positive relationship arises from a general process overlap (e.g. multivariate patterns of encoding), which is not specific to individual items but would be similarly present in other trial -to-trial pairs that do not involve the same stimulus. Another alternative is that coarser context information is reinstated instead of item-specific representations.

To test whether the positive relationship between R_1,2_ and memory was specific to intact item pairs, we ran a permutation analysis where we kept the trial of the 1^st^ presentation fixed but shuffled the activation vectors used as the 2^nd^ presentation trial (iterations = 10,000). One-sided p-values were derived by fitting a function to the null distribution via logspline density estimation to determine whether the β estimate of the intact pairs exceeded the null distribution.

To examine whether reinstatement was based on context instead of items, we estimated context-reinstatement by keeping the trial of the 2^nd^ encounter fixed and correlating it with all other images where both the 1^st^ and the 2^nd^ presentation occurred in the same sessions but not in the same runs as the intact pair (Zou et al., 2023). For example, Images 1 and 2 both occurred in Session 3 and Session 10 for their respective 1^st^ and 2^nd^ encounter. Context-reinstatement was defined as the median of all context pairs and then entered into a joint model alongside item-specific R_1,2_.

### Novelty analyses

To contrast the effects of consistency as measured by reinstatement with hippocampal differentiation, we calculated an item-wise novelty response by subtracting the single-trial β estimate of the 2^nd^ encounter from the corresponding value of the 1^st^ encounter (hereafter referred to as N_1,2_) averaged within each subfield. Positive values of this difference score indicate that activation was higher for the initial encounter. We started the examination by probing whether the hippocampus showed a novelty response comparable to that reported in previous investigations (Quent, Zhuang, et al., 2026). To do this, we contrasted the first against the second presentation for all subfields. Then, as with R _1,2_, N_1,2_ was used to predict subsequent memory performance in the subfields showing evidence for hippocampal reinstatement based on the expectation that the novelty response partly reflects the differential need for encoding, which would be reduced for the 2^nd^ encounter.

### Characterisation of novelty response and reinstatement

To further characterise the relationship between the novelty response and reinstatement, we examined whether both co-vary by predicting image-wise reinstatement based on the image-wise novelty response. One of the key advantages of NSD is the broad range of lags between presentations allowing us to conduct a final characterisation of the two neural indices: testing whether their expressions are time-limited in line with the idea that the hippocampus is only needed until a memory is consolidated (Squire & Alvarez, 1995). Thus, we used the log-transformed lags between the 1^st^ and the 2^nd^ encounter to separately predict R_1,2_ and N_1,2_.

## Results

### Behavioural results

As previously reported, NSD features a high level of data quality, including the behavioural performance over the course of the whole experiment. Nevertheless, across all participants, memory performance decreased over time as indicated by session-level averages of d-prime (Figure 1B), β = -1.44 (95 % CI [-1.91, -0.94]). Across the whole experiment, d-prime ranged from 0.89 to 1.60 with an average of 1.25 (SD = 0.27). This decline in memory performance can be partly attributed to the increasing number of trials with large lags between presentations of the same stimulus. Lags ranged from minutes to months (mean = 46.5, median = 10, SD = 64.5 days; Figure 1C) in the NSD and negatively predicted retrieval success with a large effect size. This can be illustrated by predicting retrieval success based on the lag between the 2^nd^ and the 3^rd^ encounter (Figure 1C), β = -2.86 (95 % CI [-3.84, -1.67]). Similarly, the lag between the 1^st^ and the 2^nd^ encounter negatively predicted retrieval success for the 2^nd^ presentation, β = -2.53 (95 % CI [-3.57, -1.31]). To isolate the effects of reinstatement and novelty from such lag-induced retrieval failure, lag and its interaction were always added as co-variates of no interest.

### Reinstatement of 1^st^ encounter predicts subsequent memory in CA1 and CA3

In line with our main hypothesis, R_1,2_ within CA1 robustly predicted successful memory retrieval for the 3^rd^ encounter of an image (Figure 2), β = 0.069 (95 % CI [0.004, 0.136]), illustrating that hippocampal reinstatement of the initial experience can drive successful memory formation in a simple recognition task. In additional exploratory analyses, we investigated whether other subfields similarly showed evidence of hippocampal reinstatement. In addition to CA1, we found that hippocampal reinstatement in CA3 predicted subsequent memory as well, β = 0.082 (95 % CI [0.011, 0.153]). In contrast, no robust effects were observed for any other subfield (subiculum, CA2 or DG). Given this incidental finding, we further investigated the nature of reinstatement in CA3 alongside the planned analyses for CA1.

After establishing that hippocampal reinstatement can drive recognition memory, we next wondered whether hippocampal reinstatement effects in CA1 and CA3 were restricted to R_1,2_ or whether other forms of reinstatement are similarly predictive of memory. However, in contrast to R_1,2_, no other form of reinstatement robustly predicted memory for the 3^rd^ encounter: either in CA1, R_1,3_: β = 0.0264 (95 % CI [-0.0355, 0.0914]), R_2,3_: β = 0.0424 (95 % CI [-0.0234, 0.109]), or in CA3, R_1,3_: β = 0.0172 (95 % CI [-0.0829, 0.108]), R_2,3_: β = 0.0177 (95 % CI [-0.0593, 0.0878]). Thus far, we tested whether during the 3^rd^ presentation of a scene, participants would reinstate either the initial experience or its first repetition. However, we also explored the possibility that the hippocampus might reinstate the representations of the 1^st^ and the 2^nd^ presentation simultaneously. To this end, we calculated an average pattern of the 1^st^ and the 2^nd^ presentations and correlated it with the pattern evoked by the 3^rd^ encounter (R_avg,3_). However, again neither in CA1, β = 0.0575 (95 % CI [-0.000885, 0.118]), nor in CA3, β = 0.0325 (95 % CI [-0.0851, 0.144]), was there strong evidence that this form of reinstatement predicted memory. Beneficial effects of hippocampal reinstatement were therefore mostly specific to reactivation of the initial representation during the first repetition and retrieval attempt.

Finally, we also investigated whether R_1,2_ reinstatement predicted memory for the 2^nd^ encounter itself or, in other words, whether the reinstatement effects were simply a signature of retrieval success when a scene is seen for the 2^nd^ time. However, this was neither the case for CA1, β = 0.0188 (95 % CI [-0.0368, 0.0824]), nor for CA3, β = 0.00482 (95 % CI [-0.0821, 0.095]). R_1,2_’s inability to predict memory for the 2^nd^ time suggests that reinstatement is not simply a co-product of successful retrieval but might be directly tied to re-processing a stimulus.

Thus far, we observed that hippocampal reinstatement (i.e. consistent representations) in CA1 as well as in CA3 is predictive of subsequent recognition memory. Importantly, these reinstatement effects were specific to retrieving the initial experience during its first repetition.

### Item-specificity of hippocampal reinstatement

In line with previous investigations (Ritchey et al., 2013; Zou et al., 2023), we ran a series of analyses to further characterise the nature of hippocampal reinstatement. Specifically, we investigated whether reinstatement in CA1 and CA3 was item-specific or reflected a general process overlap (i.e. multivariate patterns associated with encoding). Such process overlap would likewise be expressed in other item-to-item pairs. However, we did find that the ability of reinstatement in intact scene-pairs to predict subsequent memory exceeded that of reinstatement calculated based on permuted pairs in CA1, p = .035, as well as in CA3, p = .003, suggesting that reinstatement was indeed specific to intact pairs (Figure 3).

**Figure 3.**
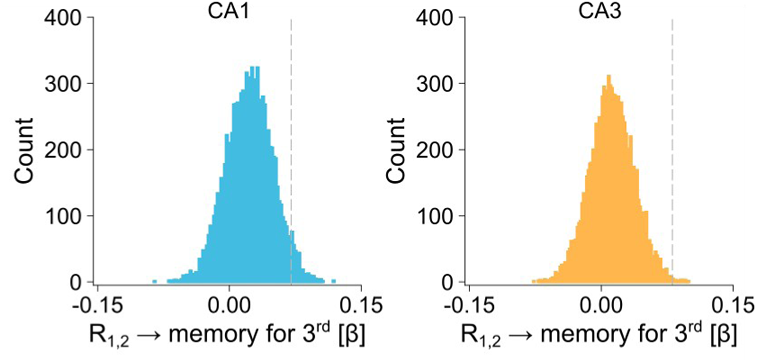
Permutation analysis of item specificity. Null distributions shown as histograms derived from permutation analyses of shuffling activation vectors (iterations = 10,000). The vertical lines represent the empirical estimates based on intact pairs.

We additionally examined whether reinstatement was driven by retrieving item-specific or more generally context-specific information. To this end, we calculated reinstatement between the 2^nd^ presentation and scenes that shared the same context (i.e. same sessions). We then entered this context-specific R_1,2_ alongside the item-specific R_1,2_ into a joint model to control for each other’s contribution to successful memory. This analysis uncovered a notable dissociation between CA1 and CA3. In CA1, item-specific reinstatement, β = 0.0612 (95 % CI [-0.00167, 0.128]), continued to show a similar effect size but could not be separated cleanly from context-specific reinstatement, β = 0.0708 (95 % CI [-0.023, 0.164]). In contrast, in CA3, there was no evidence for any effect of context-specific reinstatement, β = 0.00964 (95 % CI [-0.0756, 0.0915]), but clear evidence for item-specific reinstatement, β = 0.0807 (95 % CI [0.00539, 0.155]). We further directly contrasted R_1,2_ in CA1 versus CA3 to probe the different nature of their reinstatement. However, with the current data, we were not able to separate the effects of CA1, β = 0.0496 (95 % CI [-0.0159, 0.117]), and CA3, β = 0.0701 (95 % CI [-0.00608, 0.144]), despite again showing similar effect sizes in this joint model. Collectively, this suggests that reinstatement in CA1 does not represent a generic process overlap but instead represents reactivations of specific information. In the case of CA3, this reinstatement was exclusively specific to item-level information.

### Independent contributions of hippocampal reinstatement and novelty response

So far, our results have highlighted that consistency in the hippocampus is conducive to successful memory. However, a rich literature also suggests that the novelty response (i.e. the activation difference between presentations) supports memory encoding. We therefore first sought evidence that the hippocampus is sensitive to novelty in the NSD. We found that N_1,2_ responses were robust across all hippocampal subfields (Figure 4), with a peak in CA3, β = 0.09 (95 % CI [0.04, 0.14]), in line with previous work (Quent, Zhuang, et al., 2026) but robust in CA1, β = 0.05 (95 % CI [0.02, 0.08]), suggesting the same subfields differentiated repetitions in addition to expressing consistent representations.

**Figure 4.**
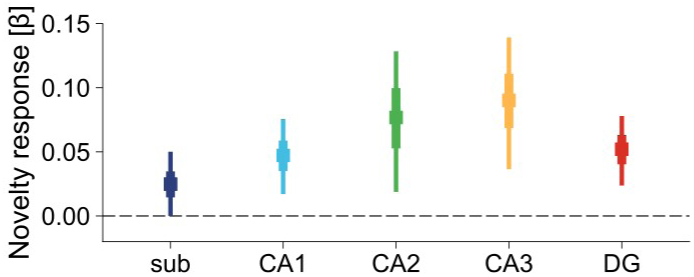
Univariate novelty response across the subfields. Posterior distributions of fixed effects of the novelty response (i.e. 1^st^ > 2^nd^) are summarised via their median (centre square) and the 66% and 95% CI.

To formally probe the novelty response’s role in memory encoding, we predicted memory for the 3^rd^ presentation following the logic of the reinstatement analyses. We found that N_1,2_ in CA1, β = 0.0988 (95 % CI [0.0232, 0.17]), but not in CA3, β = 0.00385 (95 % CI [-0.0617, 0.0708]), robustly predicted subsequent memory. This suggests that in CA1 but not in CA3, the novelty response partly reflects differential encoding activity. There was some evidence that CA3’s novelty response simply reflected retrieval success instead, as N_1,2_ predicted memory for the 2^nd^ presentation, β = 0.226 (95 % CI [0.11, 0.341]), as did the novelty response in CA1, β = 0.116 (95 % CI [0.0122, 0.214]). Furthermore, this CA3 model was the only model reported thus far where the relationship between the neural index and memory was qualified by an interaction with the log-transformed lag, β = -0.196 (95 % CI [-0.352, -0.0438]), supporting the idea that CA3’s novelty is only associated with retrieval but not with encoding. These findings corroborate the suggestion that only CA1’s novelty response contributes to memory encoding.

To ensure that these effects involving N_1,2_ indeed reflected differential hippocampal activation, we further ran an alternative model that summed activation across both initial encounters instead of subtracting. In line with a differentiation account, the summed response robustly but negatively predicted memory for the 3^rd^ encounter in CA1, β = -0.0988 (95 % CI [-0.182, -0.00111]), with no robust association found for CA3 in either direction, β = 0.0496 (95 % CI [-0.0687, 0.171]). This indicates that the novelty response in CA1 (i.e. differentiation) is the univariate signature of successful memory formation.

Finally, we tested whether hippocampal consistency (i.e. R_1,2_) and hippocampal differentiation (i.e. N_1,2_) provide independent contributions to successful memory when entered into a joint model (Figure 5). For CA1, both N_1,2_, β = 0.0985 (95 % CI [0.0248, 0.167]), as well as R_1,2_, β = 0.0663 (95 % CI [0.00263, 0.132]), continued to predict memory performance even after controlling for each other’s variance. For completeness, we also repeated this analysis for CA3 despite no evidence that the novelty response within this subfield contributed to memory encoding. As expected, N_1,2_, β = 0.00467 (95 % CI [-0.0616, 0.0721]), did not predict subsequent memory while R_1,2_ continued to, β = 0.0825 (95 % CI [0.00851, 0.153]), even after controlling for the univariate novelty response. These results are not surprising because we also found that both in CA1, β = -0.00108 (95 % CI [-0.0332, 0.0298]), and in CA3, β = -0.0364 (95 % CI [-0.0986, 0.0238]), R_1,2_ did not correlate with N_1,2_. Thus the novelty response and reinstatement are orthogonal signals, which independently contribute to memory.

**Figure 5.**
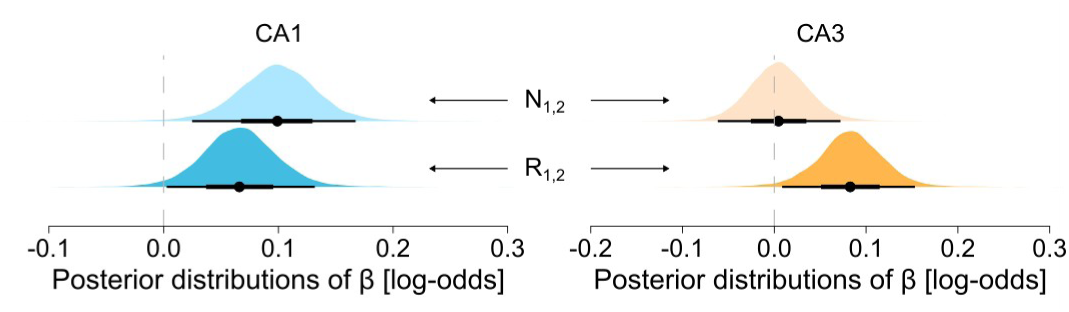
Joint models of novelty response and reinstatement. Posterior distributions of fixed effects with R1,2 and N1,2 in joint models are summarised via their median (centre dot) and the 66% and 95% CI.

### Reinstatement and novelty response do not diminish over time

After establishing that the novelty response and reinstatement represent orthogonal signals, we further explored how they themselves are influenced by the passage of time between presentations. The theory of standard system consolidation (Squire & Alvarez, 1995) posits a time-limited contribution of the hippocampus. Based on this, we would expect that reinstatement would decrease as the lag increases, especially given the large range of lags which could be up to 300 days. However, neither in CA1, β = -0.0106 (95 % CI [-0.0442, 0.0235]), nor in CA3, β = -0.0218 (95 % CI [-0.0445, 0.00145]), was this found to be the case. The same was observed for the novelty response, CA1: β = -0.0368 (95 % CI [-0.131, 0.0606]), CA3: β = -0.0716 (95 % CI [-0.188, 0.0435]). Despite all regression coefficients being negative, which indicates a reduction of the indices, there was no strong evidence that any of them changed over time.

## Discussion

Across our analyses, we found evidence that consistency as well as differentiation within the hippocampus can be conducive to successful memory. With regard to consistency, we found that hippocampal reinstatement in our primary subfield, CA1, supported memory formation. In additional exploratory analyses, we also found similarly beneficial reinstatement in CA3. No other subfield of the hippocampus expressed similar patterns. While reinstatement in CA3 was exclusively item-specific, we could not distinguish item-specific from context-specific reinstatement in CA1, possibly due to both sharing similar variance. We found evidence for beneficial differentiation, as the novelty response in CA1 contributed to memory encoding. Importantly, this contribution of hippocampal differentiation supported memory encoding independently of the extent to which the hippocampus expressed consistent representations across repetitions.

These findings thus provide evidence that the hippocampus not only flexibly shifts between integration and differentiation in response to changing goals (Brunec et al., 2020), but also expresses beneficial forms of consistency and differentiation at the same time. While previous work suggested that the cortical reinstatement but not the cortical novelty response predicts recognition memory (Ward et al., 2013), we report that novelty and reinstatement within the hippocampus can be orthogonal to each other, in particular in CA1, and simultaneously be predictive of successful memory formation.

This raises the question of what the mechanisms behind these orthogonal signatures are. The novelty response in CA1 likely reflects increased encoding activity in response to detecting new or unfamiliar information (Maass et al., 2014; Tulving et al., 1996; Tulving & Kroll, 1995) – a core feature attributed to the hippocampus (Quent et al., 2021). The 2^nd^ presentation, which relatively speaking, requires less encoding, serves as a stimulus-specific baseline for the 1^st^ presentation. Yet, this explanation does not preclude the possibility that the presentations additionally differ across other dimensions such as attention. With regard to reinstatement, we found that within the hippocampus it does not simply reflect retrieval. Instead, it is possible that the strength of reinstatement partly reflects reconsolidation (Nader & Hardt, 2009). Reconsolidation is a process during which memories enter a transient state of instability that allows updating and/or strengthening memories, in which consistency might be especially beneficial. In our case, reinstating a memory trace during the 2^nd^ encounter might have provided the opportunity for reconsolidation. Here, our results furthermore contradict the idea that the role of the hippocampus is time-limited, at least with regard to novelty and reinstatement (Squire & Alvarez, 1995), as we did not find that either varies as a function of the lag between presentations.

Our findings are also notable because the hippocampus is mainly viewed as an associative system, whose main role is to bind components of events to their spatial-temporal context (Diana et al., 2007; Eichenbaum & Cohen, 2014; Tulving, 2002; Yonelinas et al., 2019). Critically, however, both consistency and differentiation signatures contributed to recognition memory here. Hippocampal involvement in this type of memory has been intensely debated (Bowles et al., 2007; Daselaar et al., 2006; Eichenbaum et al., 2007; Merkow et al., 2015; Wixted & Squire, 2010), although there are several lines of evidence that the hippocampus is engaged in non-relational memory tasks (Kim, 2013, 2017).

Mirroring this debate, prior studies reported CA1 reinstatement (Tompary et al., 2016; Zou et al., 2023) for associative forms of memory such as paired-associate learning and temporal memory. Further demonstrating the fragility of reinstatement in neuroimaging studies that do not explicitly require associative encoding, a recent registered replication study failed to show robust effects of hippocampal reinstatement on memory, suggesting that minor changes in imaging parameters can alter the interpretation of statistical results (Yuksel et al., 2026). Despite the apparent elusiveness, we report a relationship between hippocampal reinstatement and subsequent recognition memory in CA1 as well as in CA3. One explanation for this positive result could be that we did not average across subfields with diverse mnemonic functions. Another reason could be that the NSD’s scale provides a higher degree of statistical power, which traditional memory experiments do not afford. The scale of the NSD not only provides high statistical power but also means that participants had to remember an order of magnitude more memoranda (up to 10,000 individual scene stimuli) than is the case in typical studies. It is possible that memorising many scenes taxes the hippocampal memory system more, and in that respect it is more aligned with everyday memory. Future work on hippocampal reinstatement is needed to explore the boundary conditions of its role in memory encoding, especially with regard to recognition memory.

Beyond demonstrating the importance of hippocampal reinstatement in recognition memory in general, our exploratory analyses additionally identified hippocampal reinstatement in CA3. The choice to focus on CA1 was based on the evidence from human neuroimaging studies implicating this subfield in reinstatement and novelty (Chen et al., 2015; Duncan et al., 2012; Maass et al., 2014; Tompary et al., 2016; Zou et al., 2023). That being said, theoretical models, especially those based on animal work, which afford clearer anatomical separation of the subfields, also posit a critical role of CA3 in pattern completion (Leutgeb & Leutgeb, 2007; Yassa & Stark, 2011). Our findings add nuance by revealing notable differences between the subfields. On the one hand, with respect to the representational nature, reinstatement in CA3 was purely and exclusively item-specific. In contrast, item-specific reinstatement could not be cleanly disentangled from context-specific reinstatement in CA1. This suggestive evidence is in line with previous work delineating different contributions of both subfields. For example, CA1 but not CA3 is needed to retrieve context information (Ji & Maren, 2008). In the same vein, CA1 is important for recalling gist memory, while CA3 contributes to precision (Atucha et al., 2023). Our findings thus dovetail with the view that CA1 and CA3 differ in their levels of granularity and sparsity with more sparse representations in the latter (Singh & Schapiro, 2026). On the other hand, the dissociation between CA1 and CA3 is even clearer when examining their differential roles in novelty. Here, only CA1 contributed to memory encoding, while CA3 did not despite the fact that its univariate response itself was stronger. This is reminiscent of CA1’s role in novelty detection (Quent, Zhuang, et al., 2026) and in line with previous similar findings (Maass et al., 2014). Collectively, the findings from our current work suggest that CA1 and CA3 serve complementary, yet functionally distinct, roles during memory formation.

The current study also sheds light on the discussion regarding which univariate encoding signature is indicative of successful encoding in the hippocampus. According to theories, novelty processing is intimately linked to memory encoding (Kim, 2013; Quent et al., 2021; Tulving et al., 1996; Tulving & Kroll, 1995). In other words, this perspective favours differential patterns of activation. As pointed out above, the novelty response in CA1 likely reflects differences in encoding demands. However, across the brain, there is conflicting evidence with both the novelty response (Ben-Yakov et al., 2014; Turk-Browne et al., 2006) and the summed response (Manelis et al., 2013; Wagner et al., 2000; Xue et al., 2011; Xue, Mei, et al., 2010) being associated with successful subsequent memory. Here, for CA1, we found clear evidence that the novelty response is indeed the signature of successful encoding. Summed responses were negatively associated with subsequent memory, while the novelty response was positively associated with memory both for the 2^nd^ and for the 3^rd^ encounter.

While our results together support the conclusion that hippocampal differentiation and consistency simultaneously and independently contribute to successful memory, there are also limitations to consider. First, recognition memory can be supported by recollection and/or familiarity (Yonelinas, 1994) and thus by associative (e.g. being able to retrieve the exact spatiotemporal context) as well as by non-associative retrieval. Unfortunately, the NSD paradigm did not allow us to separate these processes and investigate how reinstatement and the novelty response might support recollection and familiarity in different ways. Second, the dissociation between CA1 and CA3 in terms of the nature of reinstatement is merely suggestive in the current study but deserves attention as it is in line with the clearer dissociation in our novelty analyses and with the literature at large. However, future work is needed to further improve our understanding of the complementary roles of CA1 and CA3 in memory encoding. Finally, our attempts at understanding the different mnemonic contributions of hippocampal subfields using fMRI in humans are naturally constrained by a spatial resolution that, even at 7T, is orders of magnitude lower than what can be achieved in animals. As such, the possibility of partial-volume blurring across subfields should to be acknowledged.

Together, our results show that within the hippocampus differentiation and consistency, as measured by the novelty response and pattern reinstatement respectively, co-exist as orthogonal signals and their simultaneous expression provides independent contributions to successful memory.

## Authors contributions

J.A.Q.: Conceptualisation, Formal analysis, Writing – original draft, Writing – review & editing. M.Z.: Conceptualisation, Writing – original draft, Writing – review & editing. D.V.: Conceptualisation, Supervision, Writing – original draft, Writing – review & editing.

## Code and data availability statement

Data to reproduce the analyses in this manuscript can be accessed by following the instructions of the NSD website (https://naturalscenesdataset.org/). All code used for downloading and preliminary processing of NSD can be accessed via a separate repository (https://github.com/cognizelab/NSD_HippUnfold). Code for all subsequent analyses can be found at https://github.com/cognizelab/reinstatement_novelty containing all code and dependencies needed for replication.

## Declaration of competing interest

The authors of this article declare that they have no financial conflict of interest regarding the content of this article.

## Acknowledgement

Collection of the NSD dataset was supported by NSF IIS-1822683 and NSF IIS-1822929. D.V. discloses support for this research from the Ministry of Science and Technology of China, STI2030 – Major Projects [grant number 2022ZD0207900]. J.A.Q. received funding from the China Postdoctoral Science Foundation [grant number 2022M720818] to complete this work.

## Notes

### Competing Interest Statement

The authors have declared no competing interest.

## References

Allen, E. J., St-Yves, G., Wu, Y., Breedlove, J. L., Prince, J. S., Dowdle, L. T., Nau, M., Caron, B., Pestilli, F., Charest, I., Hutchinson, J. B., Naselaris, T., & Kay, K. (2022). A massive 7T fMRI dataset to bridge cognitive neuroscience and artificial intelligence. Nature Neuroscience, 25(1), 116–126. 10.1038/s41593-021-00962-x

Atucha, E., Ku, S.-P., Lippert, M. T., & Sauvage, M. M. (2023). Recalling gist memory depends on CA1 hippocampal neurons for lifetime retention and CA3 neurons for memory precision. Cell Reports, 42(11), 113317. 10.1016/j.celrep.2023.113317

Bakker, A., Kirwan, C. B., Miller, M., & Stark, C. E. L. (2008). Pattern Separation in the Human Hippocampal CA3 and Dentate Gyrus. Science, 319(5870), 1640–1642. 10.1126/science.1152882

Barr, D. J., Levy, R., Scheepers, C., & Tily, H. J. (2013). Random effects structure for confirmatory hypothesis testing: Keep it maximal. Journal of Memory and Language, 68(3), 255–278. 10.1016/j.jml.2012.11.001

Bein, O., & Davachi, L. (2026). Adaptive processing of predictions and prediction errors in hippocampal subregions. Philosophical Transactions B, 381(1954), 20250239. 10.1098/rstb.2025.0239

Ben-Yakov, A., Rubinson, M., & Dudai, Y. (2014). Shifting Gears in Hippocampus: Temporal Dissociation between Familiarity and Novelty Signatures in a Single Event. The Journal of Neuroscience, 34(39), 12973–12981. 10.1523/JNEUROSCI.1892-14.2014

Bowles, B., Crupi, C., Mirsattari, S. M., Pigott, S. E., Parrent, A. G., Pruessner, J. C., Yonelinas, A. P., & Köhler, S. (2007). Impaired familiarity with preserved recollection after anterior temporal-lobe resection that spares the hippocampus. Proceedings of the National Academy of Sciences of the United States of America, 104(41), 16382–16387. 10.1073/pnas.0705273104

Brunec, I. K., Robin, J., Olsen, R. K., Moscovitch, M., & Barense, M. D. (2020). Integration and differentiation of hippocampal memory traces. Neuroscience & Biobehavioral Reviews, 118, 196–208. 10.1016/j.neubiorev.2020.07.024

Bürkner, P.-C. (2017). brms: An R Package for Bayesian Multilevel Models Using Stan. Journal of Statistical Software, 80(1). 10.18637/jss.v080.i01

Chen, J., Cook, P. A., & Wagner, A. D. (2015). Prediction strength modulates responses in human area CA1 to sequence violations. Journal of Neurophysiology, 114(2), 1227–1238. 10.1152/jn.00149.2015

Daselaar, S. M., Fleck, M. S., & Cabeza, R. (2006). Triple Dissociation in the Medial Temporal Lobes: Recollection, Familiarity, and Novelty. Journal of Neurophysiology, 96(4), 1902–1911. 10.1152/jn.01029.2005

DeKraker, J., Cabalo, D. G., Royer, J., Ngo, A., Khan, A. R., Karat, B. G., Benkarim, O., Rodriguez-Cruces, R., Frauscher, B., Pana, R., Hansen, J. Y., Misic, B., Valk, S. L., Lau, J. C., Kirschner, M., Bernasconi, A., Bernasconi, N., Muenzing, S. E. A., Axer, M., … Bernhardt, B. C. (2025). HippoMaps: Multiscale cartography of human hippocampal organization. Nature Methods, 22(10), 2211–2222. 10.1038/s41592-025-02783-3

DeKraker, J., Lau, J. C., Ferko, K. M., Khan, A. R., & Köhler, S. (2020). Hippocampal subfields revealed through unfolding and unsupervised clustering of laminar and morphological features in 3D BigBrain. NeuroImage, 206, 116328. 10.1016/j.neuroimage.2019.116328

DeKraker, J., Palomero-Gallagher, N., Kedo, O., Ladbon-Bernasconi, N., Muenzing, S. E. A., Axer, M., Amunts, K., Khan, A. R., Bernhardt, B., & Evans, A. C. (2023). Evaluation of surface-based hippocampal registration using ground-truth subfield definitions. 10.7554/eLife.88404.3

Diana, R. A., Yonelinas, A. P., & Ranganath, C. (2007). Imaging recollection and familiarity in the medial temporal lobe: A three-component model. Trends in Cognitive Sciences, 11(9), 379–386. 10.1016/j.tics.2007.08.001

Dimsdale-Zucker, H. R., Ritchey, M., Ekstrom, A. D., Yonelinas, A. P., & Ranganath, C. (2018). CA1 and CA3 differentially support spontaneous retrieval of episodic contexts within human hippocampal subfields. Nature Communications, 9(1), 294–294. 10.1038/s41467-017-02752-1

Duncan, K., Ketz, N., Inati, S. J., & Davachi, L. (2012). Evidence for area CA1 as a match/mismatch detector: A high-resolution fMRI study of the human hippocampus. Hippocampus, 22(3), 389–398. 10.1002/hipo.20933

Eichenbaum, H., & Cohen, N. J. (2014). Can We Reconcile the Declarative Memory and Spatial Navigation Views on Hippocampal Function? Neuron, 83(4), 764–770. 10.1016/j.neuron.2014.07.032

Eichenbaum, H., Yonelinas, A. P., & Ranganath, C. (2007). The Medial Temporal Lobe and Recognition Memory. Annual Review of Neuroscience, 30(1), 123–152. 10.1146/annurev.neuro.30.051606.094328

Hasselmo, M. E., & Wyble, B. P. (1997). Free recall and recognition in a network model of the hippocampus: Simulating effects of scopolamine on human memory function. Behavioural Brain Research, 89(1–2), 1–34. 10.1016/S0166-4328(97)00048-X

Heinbockel, H., Wagner, A. D., & Schwabe, L. (2024). Post-retrieval stress impairs subsequent memory depending on hippocampal memory trace reinstatement during reactivation. Science Advances, 10(18), eadm7504. 10.1126/sciadv.adm7504

Horner, A. J., Bisby, J. A., Bush, D., Lin, W.-J., & Burgess, N. (2015). Evidence for holistic episodic recollection via hippocampal pattern completion. Nature Communications, 6(1), 7462. 10.1038/ncomms8462

Ji, J., & Maren, S. (2008). Differential roles for hippocampal areas CA1 and CA3 in the contextual encoding and retrieval of extinguished fear. Learning & Memory, 15(4), 244–251. 10.1101/lm.794808

Kim, H. (2013). Differential neural activity in the recognition of old versus new events: An Activation Likelihood Estimation Meta-Analysis. Human Brain Mapping, 34(4), 814–836. 10.1002/hbm.21474

Kim, H. (2017). Brain regions that show repetition suppression and enhancement: A meta-analysis of 137 neuroimaging experiments. Human Brain Mapping, 38(4), 1894–1913. 10.1002/hbm.23492

Kirchhoff, B. A., Wagner, A. D., Maril, A., & Stern, C. E. (2000). Prefrontal–Temporal Circuitry for Episodic Encoding and Subsequent Memory. The Journal of Neuroscience, 20(16), 6173 LP – 6180. 10.1523/JNEUROSCI.20-16-06173.2000

Koslov, S. R., Kable, J. W., & Foster, B. L. (2024). Dissociable contributions of the medial parietal cortex to recognition memory. *The Journal of Neuroscience*, e2220232024. 10.1523/JNEUROSCI.2220-23.2024

Kumaran, D., & Maguire, E. A. (2009). Novelty signals: A window into hippocampal information processing. Trends in Cognitive Sciences, 13(2), 47–54. 10.1016/j.tics.2008.11.004

Leutgeb, S., & Leutgeb, J. K. (2007). Pattern separation, pattern completion, and new neuronal codes within a continuous CA3 map. Learning & Memory, 14(11), 745–757. 10.1101/lm.703907

Libby, L. A., Reagh, Z. M., Bouffard, N. R., Ragland, J. D., & Ranganath, C. (2019). The Hippocampus Generalizes across Memories that Share Item and Context Information. Journal of Cognitive Neuroscience, 31(1), 24–35. 10.1162/jocn_a_01345

Lin, T.-Y., Maire, M., Belongie, S., Hays, J., Perona, P., Ramanan, D., Dollár, P., & Zitnick, C. L. (2014). Microsoft COCO: Common Objects in Context. In D. Fleet, T. Pajdla, B. Schiele, & T. Tuytelaars (Eds.), Computer Vision – ECCV 2014 (pp. 740–755). Springer International Publishing.

Maass, A., Schütze, H., Speck, O., Yonelinas, A., Tempelmann, C., Heinze, H.-J., Berron, D., Cardenas-Blanco, A., Brodersen, K. H., Enno Stephan, K., & Düzel, E. (2014). Laminar activity in the hippocampus and entorhinal cortex related to novelty and episodic encoding. Nature Communications, 5, 5547–5547. 10.1038/ncomms6547

Manelis, A., Paynter, C. A., Wheeler, M. E., & Reder, L. M. (2013). Repetition related changes in activation and functional connectivity in hippocampus predict subsequent memory. Hippocampus, 23(1), 53–65. 10.1002/hipo.22053

Merkow, M. B., Burke, J. F., & Kahana, M. J. (2015). The human hippocampus contributes to both the recollection and familiarity components of recognition memory. Proceedings of the National Academy of Sciences, 112(46), 14378–14383. 10.1073/pnas.1513145112

Nader, K., & Hardt, O. (2009). A single standard for memory: The case for reconsolidation. Nature Reviews Neuroscience, 10(3), 224 –234. 10.1038/nrn2590

Norman, K. A. (2010). How hippocampus and cortex contribute to recognition memory: Revisiting the complementary learning systems model. Hippocampus, 20(11), 1217–1227. 10.1002/hipo.20855

Prince, J. S., Charest, I., Kurzawski, J. W., Pyles, J. A., Tarr, M. J., & Kay, K. N. (2022). Improving the accuracy of single-trial fMRI response estimates using GLMsingle. eLife, 11, e77599. 10.7554/eLife.77599

Quent, J. A., Henson, R. N., & Greve, A. (2021). A predictive account of how novelty influences declarative memory. Neurobiology of Learning and Memory, 179, 107382–107382. 10.1016/j.nlm.2021.107382

Quent, J. A., Song, L., Liang, X., Su, Y., Yu, W., Wang, H., & Vatansever, D. (2026). Graded encoding of spatial novelty scales in the human brain. Nature Communications, 17(1), 303. 10.1038/s41467-025-67012-z

Quent, J. A., Zhuang, K., Liang, X., Zeng, D., Wang, Y., Feng, J., DeKraker, J., Bernhardt, B. C., & Vatansever, D. (2026). Hippocampal subfields support human novelty detection via distinct signals. bioRxiv. 10.64898/2026.09.08.748144

Ritchey, M., Wing, E. A., LaBar, K. S., & Cabeza, R. (2013). Neural similarity between encoding and retrieval is related to memory via hippocampal interactions. *Cerebral Cortex (New York*, N.Y*. :* 1991), *23*(12), 2818–2828. 10.1093/cercor/bhs258

Schlichting, M. L., Zeithamova, D., & Preston, A. R. (2014). CA1 subfield contributions to memory integration and inference. Hippocampus, 24(10), 1248–1260. 10.1002/hipo.22310

Silva, M., Wu, X., & Fuentemilla, L. (2026). Neural reactivation supports memory formation when events end. *Trends in Cognitive Sciences*, S1364661326001609. 10.1016/j.tics.2026.07.002

Singh, D., & Schapiro, A. C. (2026). Evidence for complementary learning systems within the hippocampus. Philosophical Transactions B, 381(1954), 20250243. 10.1098/rstb.2025.0243

Squire, L. R., & Alvarez, P. (1995). Retrograde amnesia and memory consolidation: A neurobiological perspective. Current Opinion in Neurobiology, 5(2), 169–177. 10.1016/0959-4388(95)80023-9

Staresina, B. P., Henson, R. N. A., Kriegeskorte, N., & Alink, A. (2012). Episodic Reinstatement in the Medial Temporal Lobe. The Journal of Neuroscience, 32(50), 18150–18156. 10.1523/JNEUROSCI.4156-12.2012

Tompary, A., Duncan, K., & Davachi, L. (2016). High-resolution investigation of memory-specific reinstatement in the hippocampus and perirhinal cortex. Hippocampus, 26(8), 995–1007. 10.1002/hipo.22582

Tulving, E. (2002). Episodic Memory: From Mind to Brain. Annual Review of Psychology, 53(1), 1–25. 10.1146/annurev.psych.53.100901.135114

Tulving, E., & Kroll, N. (1995). Novelty assessment in the brain and long-term memory encoding. Psychonomic Bulletin & Review, 2(3), 387–390. 10.3758/BF03210977

Tulving, E., Markowitsch, H. J., Craik, F. I. M., Habib, R., & Houle, S. (1996). Novelty and Familiarity Activations in PET Studies of Memory Encoding and Retrieval. Cerebral Cortex, 6(1), 71–79. 10.1093/cercor/6.1.71

Turk-Browne, N. B., Yi, D.-J., & Chun, M. M. (2006). Linking Implicit and Explicit Memory: Common Encoding Factors and Shared Representations. Neuron, 49(6), 917–927. 10.1016/j.neuron.2006.01.030

Vanasse, T. J., Boly, M., Allen, E. J., Wu, Y., Naselaris, T., Kay, K., Cirelli, C., & Tononi, G. (2022). Multiple traces and altered signal-to-noise in systems consolidation: Evidence from the 7T fMRI Natural Scenes Dataset. Proceedings of the National Academy of Sciences, 119(44), e2123426119. 10.1073/pnas.2123426119

Wagner, A. D., Maril, A., & Schacter, D. L. (2000). Interactions Between Forms of Memory: When Priming Hinders New Episodic Learning. Journal of Cognitive Neuroscience, 12(Supplement 2), 52–60. 10.1162/089892900564064

Ward, E. J., Chun, M. M., & Kuhl, B. A. (2013). Repetition Suppression and Multi-Voxel Pattern Similarity Differentially Track Implicit and Explicit Visual Memory. The Journal of Neuroscience, 33(37), 14749–14757. 10.1523/JNEUROSCI.4889-12.2013

Wing, E. A., Ritchey, M., & Cabeza, R. (2015). Reinstatement of Individual Past Events Revealed by the Similarity of Distributed Activation Patterns during Encoding and Retrieval. Journal of Cognitive Neuroscience, 27(4), 679–691. 10.1162/jocn_a_00740

Wixted, J. T., & Squire, L. R. (2010). The role of the human hippocampus in familiarity-based and recollection-based recognition memory. Behavioural Brain Research, 215(2), 197–208. 10.1016/j.bbr.2010.04.020

Xue, G., Dong, Q., Chen, C., Lu, Z., Mumford, J. A., & Poldrack, R. A. (2010). Greater Neural Pattern Similarity Across Repetitions Is Associated with Better Memory. Science, 330(6000), 97–101. 10.1126/science.1193125

Xue, G., Mei, L., Chen, C., Lu, Z. L., Poldrack, R. A., & Dong, Q. (2010). Facilitating memory for novel characters by reducing neural repetition suppression in the left fusiform cortex. PLoS ONE, 5(10), 1–10. 10.1371/journal.pone.0013204

Xue, G., Mei, L., Chen, C., Lu, Z.-L., Poldrack, R., & Dong, Q. (2011). Spaced Learning Enhances Subsequent Recognition Memory by Reducing Neural Repetition Suppression. Journal of Cognitive Neuroscience, 23(7), 1624–1633. 10.1162/jocn.2010.21532

Yassa, M. A., & Stark, C. E. L. (2011). Pattern separation in the hippocampus. Trends in Neurosciences, 34(10), 515–525. 10.1016/j.tins.2011.06.006

Yonelinas, A. P. (1994). Receiver-operating characteristics in recognition memory: Evidence for a dual-process model. *Journal of Experimental Psychology: Learning*, Memory, and Cognition, 20(6), 1341–1341.

Yonelinas, A. P., Ranganath, C., Ekstrom, A. D., & Wiltgen, B. J. (2019). A contextual binding theory of episodic memory: Systems consolidation reconsidered. Nature Reviews Neuroscience, 20(6), 364–375. 10.1038/s41583-019-0150-4

Yuksel, E., Shafer, E. S., Netto, M., & Diana, R. A. (2026). Replicability of representational similarity and its role in successful memory retrieval. Imaging Neuroscience, 4, IMAG.a.1214. 10.1162/IMAG.a.1214

Zou, F., Wanjia, G., Allen, E. J., Wu, Y., Charest, I., Naselaris, T., Kay, K., Kuhl, B. A., Hutchinson, J. B., & DuBrow, S. (2023). Re-expression of CA1 and entorhinal activity patterns preserves temporal context memory at long timescales. Nature Communications, 14(1), 4350. 10.1038/s41467-023-40100-8

